# Deconstructing a visual signal: the role of motion and colour in predator deterrence

**DOI:** 10.1101/2022.03.01.482534

**Authors:** Dinesh Rao, Skye M. Long, Horacio Tapia-McClung, Kevin Salgado-Espinosa, Ajay Narendra, Samuel Aguilar-Arguello, Luis Robledo-Ospina, Dulce Rodriguez-Morales, Elizabeth M. Jakob

## Abstract

Visual animal communication, whether to the same species or to other species, is largely conducted through dynamic and colourful signals. For a signal to be effective, the signaller must capture and retain the attention of the receiver. Signal efficacy is also dependent on the sensory limitations of the receiver. However, most signalling studies consider movement and colour separately, resulting in a partial understanding of the signal in question. We explored the structure and function of predator-prey signalling in the jumping spider-tephritid fly system, where the prey performs a wing waving display that deters an attack from the predator. Using a custom-built spider retinal tracker combined with visual modelling, and behavioural assays, we studied the effect of fly wing movement and colour on the jumping spider’s visual system. We show that jumping spiders track their prey less effectively during wing display and this can be attributed to a series of fluctuations in chromatic and achromatic contrasts arising from the wing movements. These results suggest that displaying flies deter spider attacks by manipulating the movement biases of the spider’s visual system. Our results emphasise the importance of receiver attention on the evolution of interspecific communication.

## Introduction

For visual communication to be effective, a signal must attract and hold the attention of the targeted receiver, and be interpreted accurately. A number of constraints modulate the efficacy of the signal, such as the receiver’s visual capabilities and sensory biases, the medium of transmission and distance between individuals. (see (Rosenthal, 2007) for a review). The incredible variety of movements and colours in peacock spiders (Otto and Hill, 2017) or in the birds of paradise (Ligon et al., 2018) are a testament to the effect of selection in the generation of such displays. The key to triggering the receiver’s attention is a combination of display and colour (Rosenthal, 2007), but this combination has rarely been studied together.

Visual signals have led to the evolution of different behaviours in various animals in the context of sexual selection and prey attraction, among others. In some cases, prey species find it useful to attract and hold the attention of their potential predators either to advertise their toxicity (aposematism) or to distract predators (Ruxton et al., 2018). This is counterintuitive because prey signalling to the predators is inherently risky. However, prey signalling to predators may reduce the likelihood of a successful attack, either before the attack is launched or during the attack (Ruxton et al., 2018). Prey may also transmit information regarding their body condition (Caro, 1995) or seek to deceive the predator by adopting exaggerated postures that modify their appearance (Brodie, 1977). Signals that are used to deter attacks have been broadly characterised as pursuit deterrence signals (Hasson, 1991). Despite a longstanding interest in the function of these signals, there have been few empirical tests, and fewer still using ecologically relevant predators (Ruxton et al., 2018).

For pursuit deterrent signals to be effective, the movement, form, and colouration of the body part must be conspicuous to the predator’s visual system. Conspicuousness can be enhanced if the prey has structural colours produced by interference reflection, where visibility is influenced by the angle of light and movement (Kelley et al., 2019; Parker, 1998; Stuart-Fox et al., 2020), but see (Kjernsmo et al., 2020) for a counter example of structural colours used as camouflage. Wing interference colours (hereafter WIC, see (Shevtsova et al., 2011))) and wing specularity (i.e., specular reflectance or wing specularity) generated by the transparent part of insect wings may have an anti-predatory function (similar to that seen in the metallic colours of simulated greenbottle flies (Pike, 2015)), or constitute a signal for conspecific communication (Eichorn et al., 2017; Hawkes et al., 2019; Schultz and Fincke, 2009).

We explored the structure and function of predator-prey signalling in the jumping spider - tephritid fly system, where the fly performs a wing waving display that deters an attack from the spider. It has been shown in several studies (Greene et al., 1987; Hasson, 1995; Mather and Roitberg, 1987) that wing displays from tephritid flies deter attacks, but attention has usually been focussed on the pigmented fraction of the fly wings (e.g., (Rao and Díaz-Fleischer, 2012)), and there is no information about the effect of fly motion with respect to jumping spider vision.

Jumping spider vision has been reviewed in detail (Harland et al., 2012; Hill, 2022; Morehouse, 2020) and we include only relevant highlights here. Jumping spiders have an unusual distributed visual system with four pairs of eyes, two pairs of which are forward facing. Of these forward-facing eyes, the anterior lateral eyes have larger retinas that do not move and serve as excellent motion detectors. The principal eyes have small, boomerang-shaped retinas situated at the proximal end of moveable eye tubes inside the spider’s cephalothorax. The eyes tubes can move to direct the gaze of the boomerang-shaped retinas to different areas of the visual field. The anterior lateral and principal eyes work in close collaboration, as the anterior lateral eyes are necessary to direct the gaze of the principal eyes (Jakob et al. 2018). When detecting a stimulus of interest, a spider pivots its body to bring the stimulus into the field of view of the forward-facing eyes. The principal eyes can then move to track a moving stimulus or investigate a stimulus while the body remains motionless (Land, 1969), a potentially useful trait for a visually hunting predator.

We hypothesised that the pursuit deterrent effect of a displaying fly can be attributed to changes in the spider’s visual attention. To address this, we (a) analysed the change in body orientation of untethered spiders in an arena as they encountered displaying and non-displaying flies; (b) used a custom-built eyetracker to analyse the gaze direction of a tethered spider’s principal eyes as they observed videos of displaying, moving and non-moving flies; and (c) used multispectral digital photography, visual modelling of the acuity and colour perception of the spider, and empirical behavioural assays to evaluate the efficacy of chromatic and achromatic cues in deterring attacks under different light conditions.

## Methods

### Study species

We used adult female *Phidippus audax* (Araneae: Salticidae) spiders and the Mexican fruit fly (*Anastrepha ludens*; Diptera: Tephritidae). Both species are known to co-occur in citrus orchards in Mexico (Gonzalez-Lopez et al., 2015). The fruit flies frequently congregate in the leaves of citrus plants and are known to employ a lekking mating system where males defend non resource territories (Aluja et al., 2006). Female flies defend oviposition sites. Defence against conspecifics and heterospecifics is mainly with wing waving displays termed as supination, where the fly makes semi-circular loops while waving its wings in a synchronous or asynchronous manner as it approaches the target (see Fig. 1A). This display also deters attacks from jumping spiders (Rao and Díaz-Fleischer, 2012). The supination display is triggered by movement of the opponent and is similar irrespective of the identity of the opponent (Aguilar-Argüello et al., 2015).

**Figure 1.**
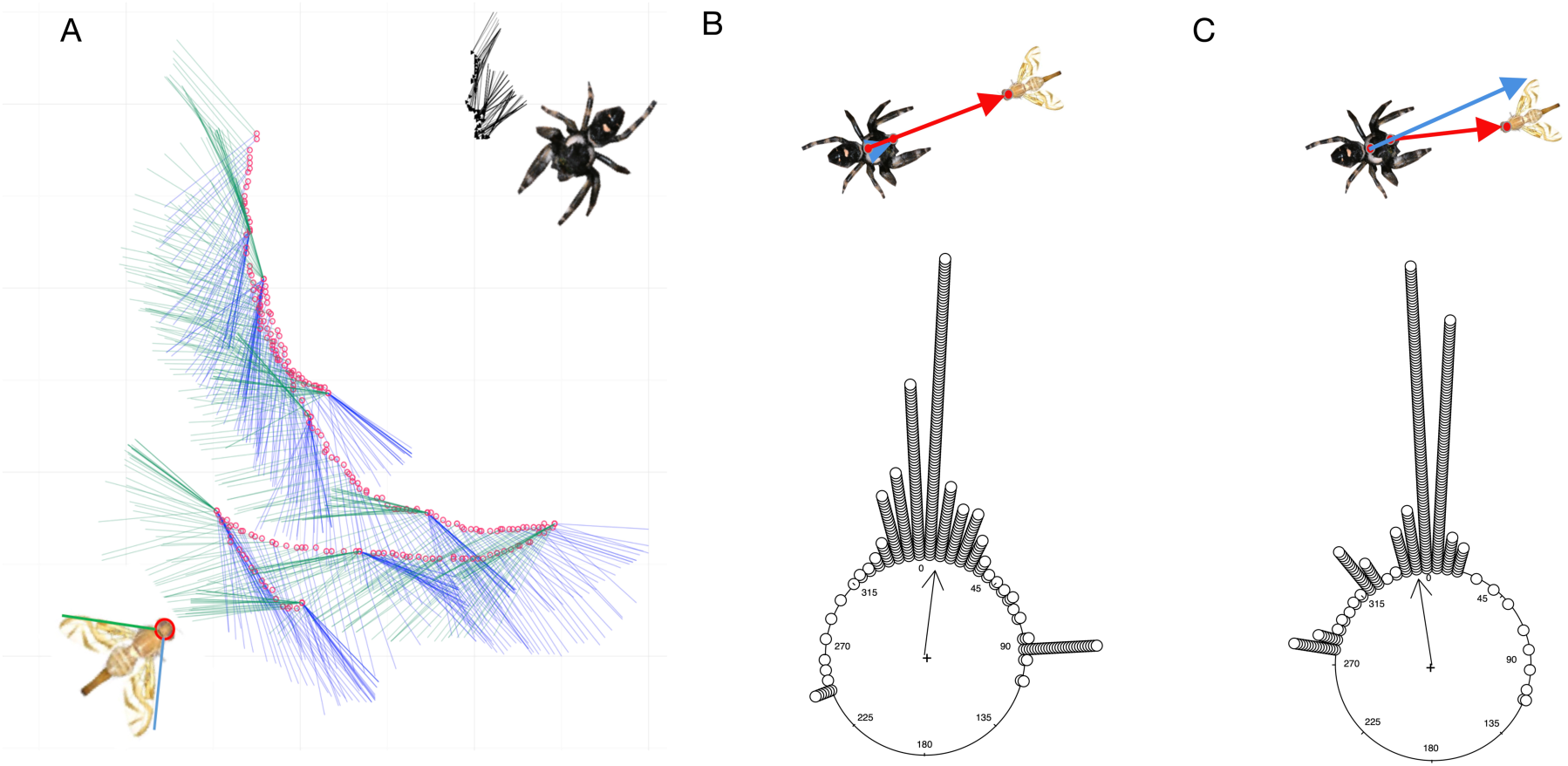
Head orientation of the spider (*Phidippus audax*) towards displaying and non-displaying Mexican fruit fly (*Anastrepha ludens*). (A) Supination display trajectory of the fly against a jumping spider predator. The red circles represent the head of the fly, the blue and green lines represent the wings. When wings are sustained in the same position, they appear as darker lines. The arrows show the body axis (head - tip of the abdomen) of the spider relative to the displaying fly. Note that the fly approaches the spider in semi looping movements during the display. Gaze direction analysis of the principal eyes of the jumping spider when facing a non-displaying (B) and a displaying fly (C). 0 degree implies that the spider gazes were aligned with the head of the fly (i.e., blue and red lines align, see inset 1B). Deviation from 0 degrees implies that spider gazes were focused away from the fly head (i.e., blue and red lines diverge in inset 1C). Display group: n=342 observations, 9 animals; Mean vector µ = 351.29°; Length of mean vector r = 0.898; circular SD: 26.51°. Non-Display group: n=342 observations, 10 animals; Mean vector µ = 9.054°; Length of mean vector r = 0.837; circular SD: 34.161°.

### Experimental design

Spiders were collected from abandoned farms around Xalapa, Mexico and kept under environmentally enriched conditions (as in (Carducci and Jakob, 2000) in the arthropod laboratory at INBIOTECA, Universidad Veracruzana. Flies were obtained as pupae from the MoscaFrut facility in Metapa de Dominguez, Chiapas, Mexico and were reared in cages. They were fed and given water *ad libitum* upon emergence. Predator-prey interactions were carried out in the laboratory under natural light conditions by placing a spider and a fly in a glass petri dish (10 cm diameter) arena, with an opaque partition separating the two. They acclimated for 1 minute and the partition was removed for the experiment to begin. The interaction was filmed from above at 25 fps at a resolution of 1920 × 1080 with a SONY HDR-PJ790V (Sony Inc.) video camera.

#### 1. Spider orientation to displaying flies in an arena

From the videos, we determined the orientation of the cephalothorax of the spiders relative to the fly by carrying out a frame-by frame analysis using a custom MATLAB program (courtesy: Jan Hemmi and Robert Parker; MathWorks, Natick, MA, USA). We manually digitised the anterior (between the spider’s principal eyes) and posterior position of the spider cephalothorax and generated x,y, coordinates to determine the approximate gaze direction of the spider’s forward-facing principal eyes. Gaze direction is a useful measure of an animal’s selective attention (Winsor et al., 2021). We did not track the position of the abdomen. We determined the position of the fly by tracking position of the center of the head. We note here that cephalothorax movement is a rough proxy of retinal movements, since jumping spider retinae have been shown to have an angular travel of up to 50° (Land, 1969; Morehouse, 2020) and it was not possible to control for movement of the eye tubes of the spiders. Nevertheless, a field of view of ± 10° from the midpoint of the sightline, i.e., a line joining the centre of the principal eyes and the back of the cephalothorax has been used in other studies (Rößler et al., 2021).

We separated videos into two categories: (a) where the fly performed a display (n = 9, S1), and (b) where there was no display (n = 10). For an illustration of a typical display, see Fig. 1A. We analysed these behaviours only when the fly and spider were subjectively observed to be oriented towards each other. To determine the gaze direction of spiders relative to the fly (Fig 1B,C inset), we calculated the angle formed by the body axis of the spider (i.e., the line connecting the anterior and posterior position of the cephalothorax of the spider) and the angle formed between the anterior position of the cephalothorax of the spider and the head of the fly. When the spider’s gaze was directed to the head of the fly, the angle recorded would be 0°. We determined this retinotopic view of spiders to displaying and non-displaying flies. Since the gaze duration varied between individuals, we used the first 38 frames of all spiders to determine the retinotopic views. From this, we determined the mean heading direction (ø), length of the mean vector (r) and circular Standard Deviation in Oriana v 4.0 (Kovach Computing Services, UK). We compared the retinal position of the spider to displaying and non-displaying flies using a Watson-Williams F test.

#### 2. Retinal tracking precision of displaying flies

In this experiment, we tracked the retinal movement of tethered spiders as they viewed a video of a fly. *Phidippus audax* spiders were collected as penultimates or adults from old fields in western Massachusetts, USA and housed in enriched environment cages at the University of Massachusetts Amherst. Spiders were fed crickets (*Acheta domesticus*) and had access to water *ad libitum*. Adult female spiders were randomly chosen from the lab population. To observe the movement of the retina of the two principal eyes (see Fig. 2A), we used a custom-built spider retina tracking apparatus (hereafter, ‘eyetracker’) at the University of Massachusetts Amherst. The eyetracker and other experimental design details are described in detail elsewhere (Canavesi et al., 2011; Jakob et al., 2018)).We provide a brief summary of the setup here.

**Figure 2.**
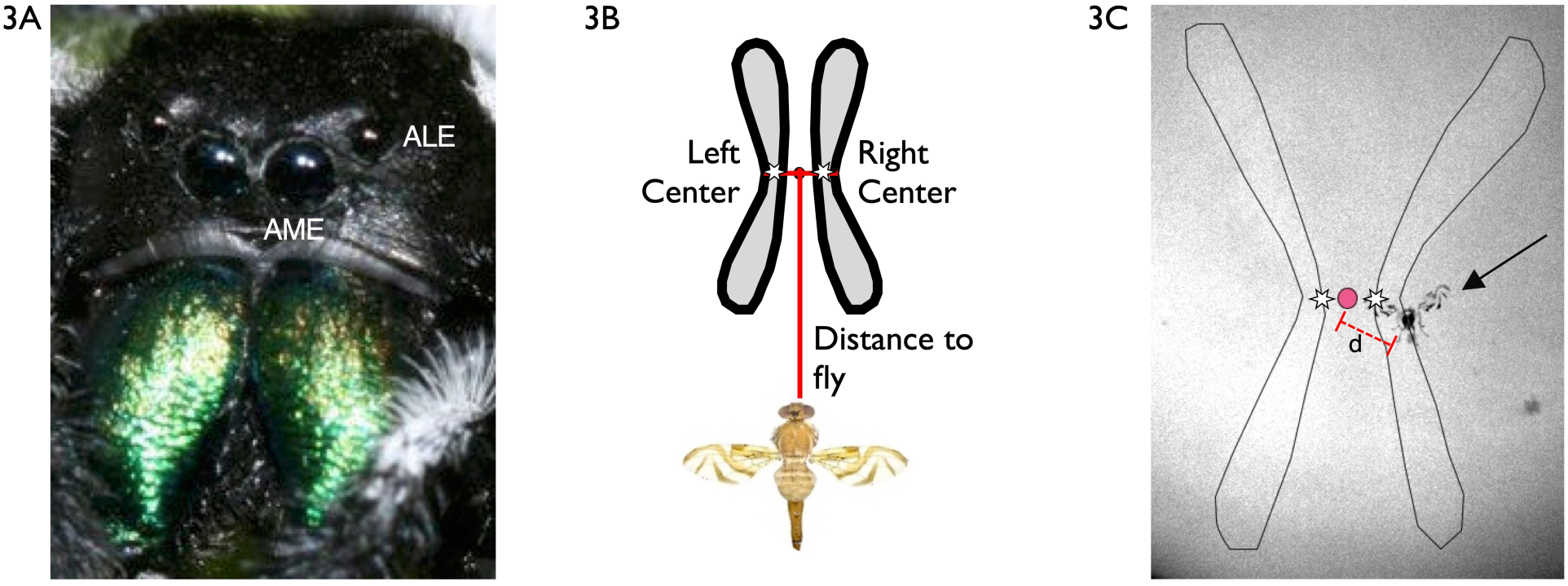
The principal eyes and retinae of the jumping spider *Phidippus audax*. (A) Frontal view of jumping spider eyes. (B) Schematic representation of the retinae of the principal eyes (i.e., Anterior Median Eyes (AME)) of the spider and (C) a frame grab showing the retinae and stimulus (i.e., a video of a fly (black arrow)). We used the midpoint (red dot) between the centre of the left and right retinae as a proxy for overall retinal movement. The distance between the midpoint and the head of the fly was used in further analysis. Note that the video of the fly was superimposed onto the video recording of retinal response to the stimulus for analysis. See text for details.

Spiders were tethered to a plastic micro-brush by a warmed wax mixture (beeswax:resin :: 1:1 mixture) applied to their cephalothorax. Spiders were aligned in the eyetracker using a manual 5-way positioner (Thor Labs) and retinae were illuminated by an IR light source (Thorlabs IR 850 nm Mounted High-Power LED). A EO-1312M CMOS monochrome USB camera recorded the motion of the retinae. The spiders viewed video stimuli projected onto a Roscoclux frosted diffusion filter (Gel/116, transmission 9%) by an Aaxa P4x Pico Projector. It has long been known that jumping spiders respond to videos with no UV information (Clark and Uetz 1990), as is the case for *P. audax* in particular (Bednarski et al. 2012, Jakob et al. 2018, Bruce et al. 2021). Proper alignment of the spider in the eyetracker was confirmed by using a calibration program written in Processing software (v2.2) and calibration videos were also used to overlay the stimulus video with the retinal position video for analysis (Jakob et al. 2018; Bruce et al. 2021). The stimulus video and the video of the retinal movements were displayed on the same computer monitor and recorded using the screen capture software Debut (v3.07).

Spiders were exposed to three types of stimulus videos: a still fly (n = 14), a moving fly (i.e., non-displaying fly that was walking; n = 15) and a displaying fly (n = 14). We used three different exemplars of the same behaviour of each stimulus treatment. The fly videos were filmed from the front (i.e., with the fly facing the camera; See S1) for the still and displaying treatments and from the side for the movement treatment. All videos were filmed with a uniform background. Each spider was presented all three fly videos in a randomised order. Spiders were exposed a white screen for 1 minute followed by 3 minutes of a stimulus video, with approximately 5 minutes of blank screen between treatments in order to allow the spider to rest.

For analysis, videos of the stimuli were overlaid on the videos of retinal movement using Final Cut Pro (Apple Inc, Cupertino). These composite videos were exported to Compressor (v. 4) and then saved as image sequences (jpg files) at 10 fps (see S2 for a sample video). Using the image sequences, we tracked the x,y position of the centres of the left and right retinae and the head of the fly (the stimulus) using the MTrackJ plugin for ImageJ ((Meijering et al., 2012), Fig. 2 B,C). We chose the head of the fly as the target since previous studies have shown that jumping spiders target the head in prey (Bartos and Minias, 2016). Only the x coordinates (i.e., horizontal displacement) were used for subsequent analysis since there is little or no vertical displacement of the fly in the videos as well as under natural conditons (see supplementary material S3 for analysis of vertical displacement).

The movements of the two principal-eye retinas of *P. audax* are highly synchronized and spiders direct the high-acuity ‘elbow’ of the boomerangs toward areas of interest (Jakob et al., 2018), so we used the midpoint of the distance between the boomerang centers as a proxy for retinal movement. The midpoint was calculated as the centre of the Euclidian distance between the centres of the two retinas.

##### 2.1 Analysis of retinal positions

We used time series analysis to analyse the data by determining the distance (i.e., separation in pixels) between stimulus and retina coordinates at each time period (i.e., video frame, 10 fps). Distances were normalised to facilitate comparison of the curves across videos. A distance value of 0 implies that the spider retinae were perfectly tracking the fly’s head. Data from pooled according to treatments. Since the order of stimulus videos were randomised, there is very little probability that any two spiders would view the fly in the same position at exactly the same time. Missing data (due to noise in video of retinae) in time and retinal position were dealt with by a 3^rd^ order Hermite interpolation. We then compared the frequency distributions of distances of all spiders between the three treatments (i.e., Display vs Moving; Display vs Still and Moving vs Still) with a Kolmogorov Smirnov test (this tests the hypothesis whether the two distributions are drawn from the same population) using Mathematica (v. 11). One outlier in the still treatment was removed due to a lack of sufficient trackable data.

##### 2.2 Optical Flow

To quantify and visualise the motion of components of the fly display and movement, we used an optical flow algorithm using the *ImageDisplacements* function in Mathematica to generate a dense motion field. This process compares horizontal and vertical displacement of individual pixels across consecutive frames of a short representative video and computes the magnitude and direction of pixel displacement (Raudies, 2013). Subsequently, these displacements were used to calculate the optical flow vector field. We then used the mean vector to present the results in a composite image for a displaying fly and a moving fly.

#### 3. Wing Interference Colouration

##### 3.1 Image acquisition and analysis

In this experiment, we simulated the appearance of the fly wing in different angles from the perspective of a jumping spider using psychophysical visual modelling techniques. We took photos of detached fly wings at different opening angles to measure colour variation during the display behaviour (termed as supination). We staged the opening angles with a wing attached to an entomological pin such that the plane of the wing was perpendicular to the ground in order to simulate different wing angles during a display. All other components of the wing position (i.e., yaw, pitch) were kept constant. We photographed the following wing opening angles: 90°, 100°, 110°, 120°, 130°, 140° and 150° against a standard grey background. We measured the relative area of the pigmented part of the wing, the wing specularity and the WIC at different opening angles in ImageJ. All photos were taken with an Olympus E-PM2 camera, modified to capture the full spectrum of light (UV-IR; but we used filters to restrict the wavelength to UV and visible light; Lifepixel, Inc.) and a EL-Nikkor 80 mm f/5.6 UV transmitting lens.

We took two photos at each angle in the ultraviolet (UV) and visible spectra (∼300 to 400 nm and ∼400 to 700 nm, respectively) of both wing and head of *A. ludens* from a front view with an Olympus Pen E-PM2 camera (converted to full-spectrum, Lifepixel.com) with a UV transmitting EL-Nikkor 80 mm f/5.6 lens attached. The ultraviolet photo was created by using a Baader UV pass and infrared (IR)/visible blocking filter, transmitting from ∼300–400 nm, and the visible spectrum photo, using a UV/IR blocking filter, transmitting between ∼400 and 700 nm. Those sets were combined for each angle using the Multispectral Image Calibration and Analysis (MICA) plugin (Troscianko and Stevens, 2015) to create a multispectral image that is a stack of images corresponding to different parts of the spectrum: ultraviolet (UV), short wavelength (SW), medium wavelength (MW), and long wavelength (LW). All the photos were taken in laboratory conditions with an Iwasaki EYE Color arc lamp (70 W 1.0 A; Venture Lighting Europe Ltd., Hertfordshire, UK) as light source. The lamp was modified by manually removing the integrated UV filter, which allows the emission of light in the UV spectrum. We did not use a diffusion filter for the lamp.

The multispectral images, with reflectance values in each channel, were transformed to the predicted photoreceptor responses (quantum catch values) for *P. audax* vision using the MICA plugin. To do this, we modelled the photoreceptors UV and MW for the spider visual system using the standard D65 illuminant, and the camera and spider spectral sensitivities. The spectral sensitivity of the camera was calculated previously (sensitivities peaks were UV 369 nm, SW 477 nm, MW 556 nm, and LW 596 nm, see (Robledo-Ospina et al., 2017)), while for *P. audax* sensitivity, we used previously reported values for the genus *Phidippus* (Peaslee and Wilson, 1989; Voe, 1975). The regression model fitted well with the photoreceptor-mapping model from camera sensitivities (R^2^ = 0.999 for both spider photoreceptors).

##### 3.2 Spatial resolution

The spatial perception linked to the visual acuity of the observer is one of the three fundamental parameters of animal vision, together with spectral sensitivity and temporal resolution (Caves et al., 2018). We simulated the resolution (acuity) of the scene viewed by the spider at different distances from the fly (2, 4, and 8 cm; these distances were selected based on previous experiments in this system (Rao and Díaz-Fleischer, 2012)). We based our analysis on the AcuityView package for R (Caves and Johnsen, 2017) but calculations were made with a custom code in the MICA toolbox, which allowed the use of quantum catch images for this simulation. We created a pseudo-colour image, which is a subset of the spider vision channels without transformation, arranged into an RGB stack, and in this case, the channel MW is shown as green and the UV as red (Fig. 5B). Details of the minimum resolvable angle calculations are presented in Supplementary material S6.

##### 3.3 Achromatic and chromatic contrasts

The details of how photoreceptors encode achromatic and colour information vary among species. However, there is evidence (summarized in (Cronin et al., 2014)) that in dichromatic animals and many arthropods (bees, flies, butterflies), the longer-wavelength sensitivity, usually associated with the MW, provides luminance information. In *Phidippus* spiders, the MW cells were the most commonly encountered type in the eye (Voe, 1975). This high proportion may support the idea that they play an essential role in the achromatic/luminance spatial vision (Cronin et al., 2014). The achromatic contrast of the wing against the standard grey background was estimated as the difference between the quantum catch value of the wing and head, divided by the sum of both values, which is also known as Michelson contrast (Olsson et al., 2018). Thus, positive values indicate that the wing is perceived as brighter than the head.

We were unable to use the traditional models for signal processing in the chromatic contrast modelling (e.g., (Vorobyev and Osorio, 1998)) due to a lack of information about either noise in each photoreceptor type or the subsequent neural processing of colour stimuli in *P. audax*. Hence, we estimated the perceived difference in spectral quality between head and wings independent of the brightness, as the Euclidean distance between those two colour signals in a triangular chromaticity space (following (Pike, 2012)), where the greater the spatial distance indicates a greater perceptual distance between the colours (Endler and Mielke, 2005). We normalized the quantum-catch values of both head and wings spectra for each photoreceptor (UV and MW) by dividing them by the sum of the quantum catches over both photoreceptor types.

##### 3.4 Behavioural assays

We manipulated the light environment in order to generate different wing appearances from the perspective of the spider. We used an UV-transmitting Teflon sheet as a light diffuser to enhance WICs.

Four light conditions were used for the behavioural assays, taken in a dark room with artificial full spectrum light (Iwasaki Eye Color Arc, Venture Lighting Europe Ltd. Hertfordshire, UK). The treatments were chosen to enhance different components of the appearance of the wing and are as follows:

1. **Control (to enhance perception of pigmented part of the wings)**: White background, direct light (i.e., no diffuser), light bulb without ultraviolet filter;
2. **Specular reflection (to enhance perception of glossy or shiny part of the wings)**: Black background, direct light, light bulb without ultraviolet filter;
3. **Wing Interference Colours (WIC) (to enhance perception of wing interference colours)**: Black background, diffuser between the light source and the wings, light bulb without ultraviolet filter;
4. **WIC without ultraviolet (to test whether UV plays a significant role in perception of wing interference colours)**: Black background, diffuser, light bulb with ultraviolet filter

In each of these four light conditions, we recorded videos of encounters (n = 15 for each treatment) between *P. audax* and *A. ludens* inside a petri dish (15 cm diameter). Before the experiment began, both the spider and the fly were placed inside the petri dish with an opaque cardboard partition (15 cm in length) in the middle for acclimatisation for a minute. We then removed the partition and we recorded the interaction for three minutes or until the spider and the fly made contact in the first interaction. We only used flies that performed the supination behaviour since previous experiments have shown that non-displaying flies are likely to be attacked at higher rates than displaying flies (Rao and Díaz-Fleischer, 2012). We quantified the number of attacks.

## Results

### Spider orientation

In arena trials, the spiders’ anterior eyes were more likely to be oriented directly at the head of non-displaying flies compared to displaying flies (F=58.49, P<0.001, Watson-Williams F-test; Fig. 1B, C).

### Retinal tracking precision

In the eyetracker, the retinas of the principal eyes of the spiders were directed more closely at the heads in videos of moving flies compared to spiders that viewed videos of nonmoving flies (Fig. 3A). Spiders facing displaying flies tracked the fly heads with the least precision (Fig. 3A). All distributions were significantly different from one another (Kolmogorov Smirnov test: Display vs Moving, D = 0.098, p < 0.01; Display vs Still, D = 0.085, p < 0.05; Moving vs Still, D = 0.072, p < 0.05; Fig. 3B).

**Figure 3.**
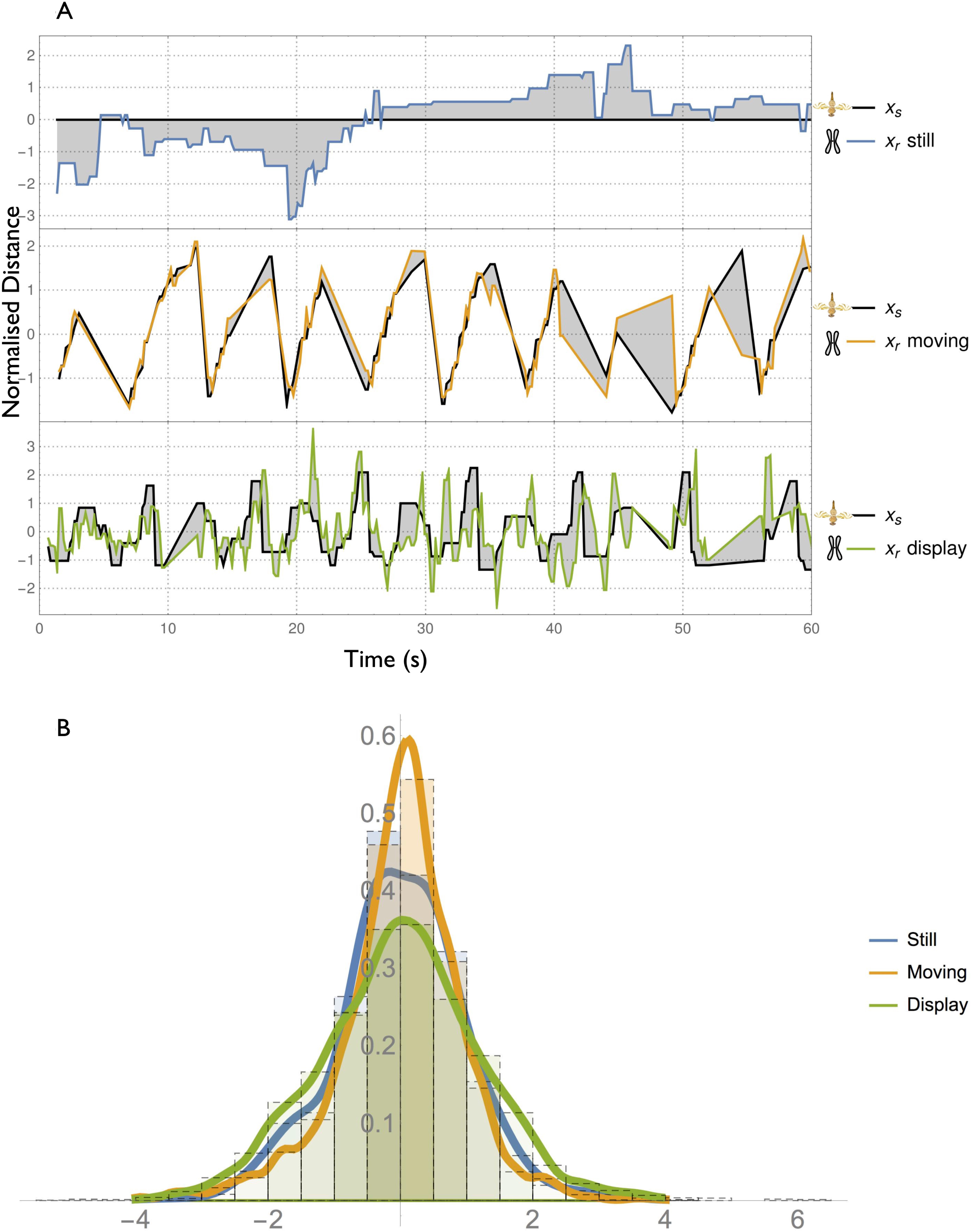
Time series analysis of the horizontal movement of stimulus and retinal response. (A) We tracked the x positions of retinae and stimulus in three treatments 1. Still fly (blue line), 2. Moving fly (orange line), and 3. Displaying fly (green line). The changes in horizontal movement over time were then normalised to have a mean of 0 and a standard deviation of 1 to facilitate comparisons between stimuli and response. The distance between the two time-series curves (shaded areas) was considered for further analysis. Perfect tracking of the stimulus would result in precisely overlapping curves. Note this is a sample figure showing responses for one spider. (B) We summarised the frequency distribution of the difference in x positions (curve separation) between the curves for the three treatments for all spiders. A data point at 0 on the x-axis implies that the stimulus and response curves coincided at that point. Retinae that tracked the stimulus better over the time period sampled would show a higher peak at 0. Moving flies were better tracked; still flies were intermediate, and displaying flies were least effectively tracked.

### Optical flow

The different motion components (i.e., a vector flow field) of a displaying fly were substantially different between a displaying fly and moving fly (Fig. 4). In general, a displaying fly produced motion in different directions and magnitudes (Fig. 4A) whereas a moving fly produced motion in one direction and at lower magnitudes (Fig. 4B). With this analysis, we noted the relative stillness of the fly’s head with respect to the wings and the ovipositor.

**Figure 4.**
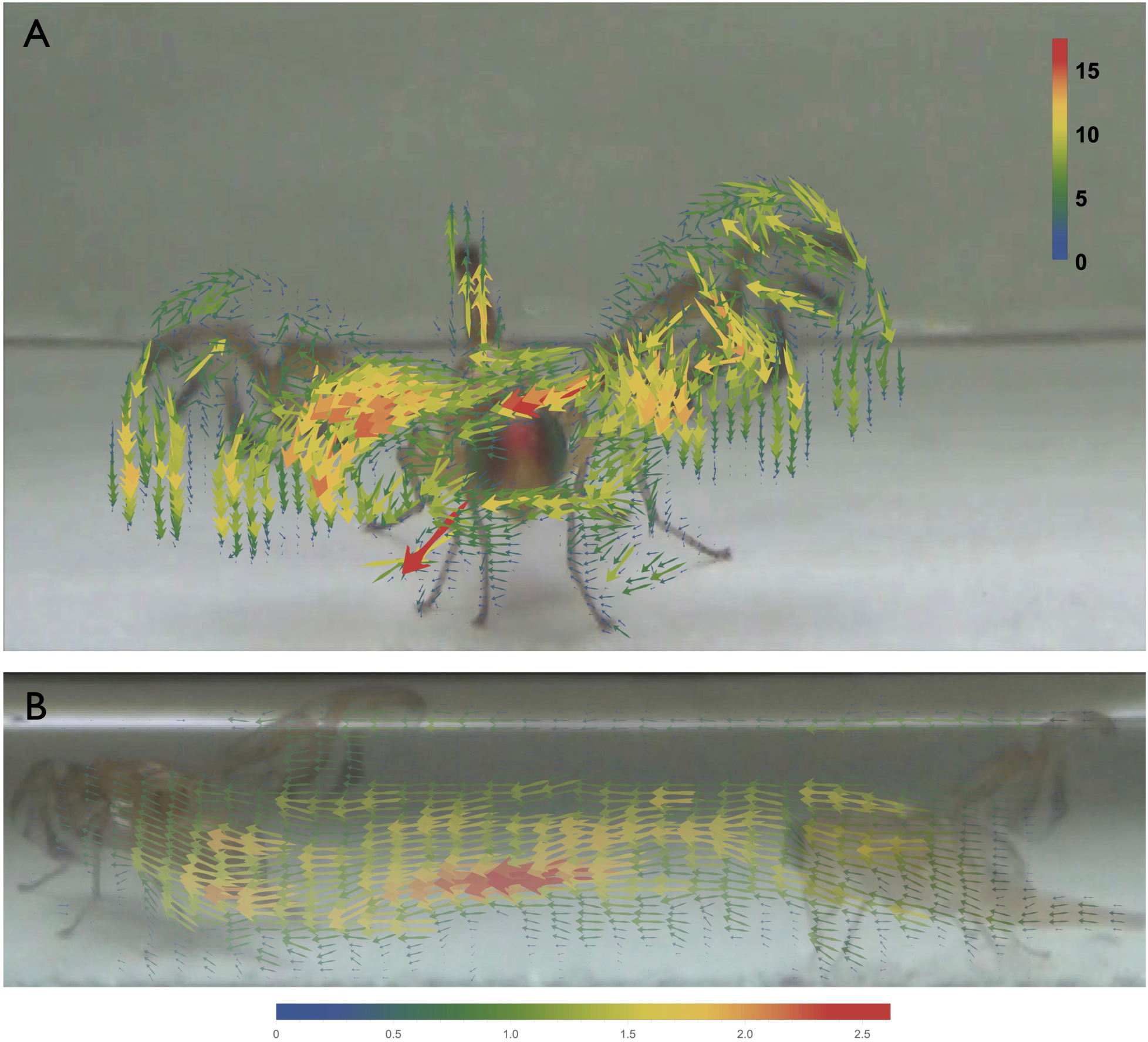
Optical flow vector fields of a (A) displaying fly and (B) a moving fly. The arrows represent direction of the pixel displacements and colour of the arrows are coded according to the mean of the vectors (i.e., magnitude of displacement).

#### Wing Interference Colouration

##### Visual acuity and modelling

We created a composite image to show the different visual components of the fly wing (Fig. 5A). According to our simulation of the fly wing appearance (Fig. 5B), spiders were likely to detect the wing specularity and bands when at a close distance and when the wing was more open (150°, Fig. 5C). The colour information produced by the transparent portion of the wing was reduced to green and UV for this species, with information from the red channel unlikely to be detected as red. At greater distances (∼8 cm), the overall appearance of the fly may be detected but details were not conspicuous. The percentage of wing area covered by WIC, UV and Specularity in general followed a hump-shaped pattern (Fig. 6A). The data was best explained by a polynomial fit to WIC (R^2^ = 0.82, F = 9.61, p < 0.05), Specularity (R^2^ = 0.71, F = 5.08, p < 0.05), UV (R^2^ = 0.91, F = 19.54, p < 0.0001) and pigment (R^2^ = 0.92, F = 26.53, p < 0.005). The significant fit suggests that information from these components is nonlinear in nature.

**Figure 5.**
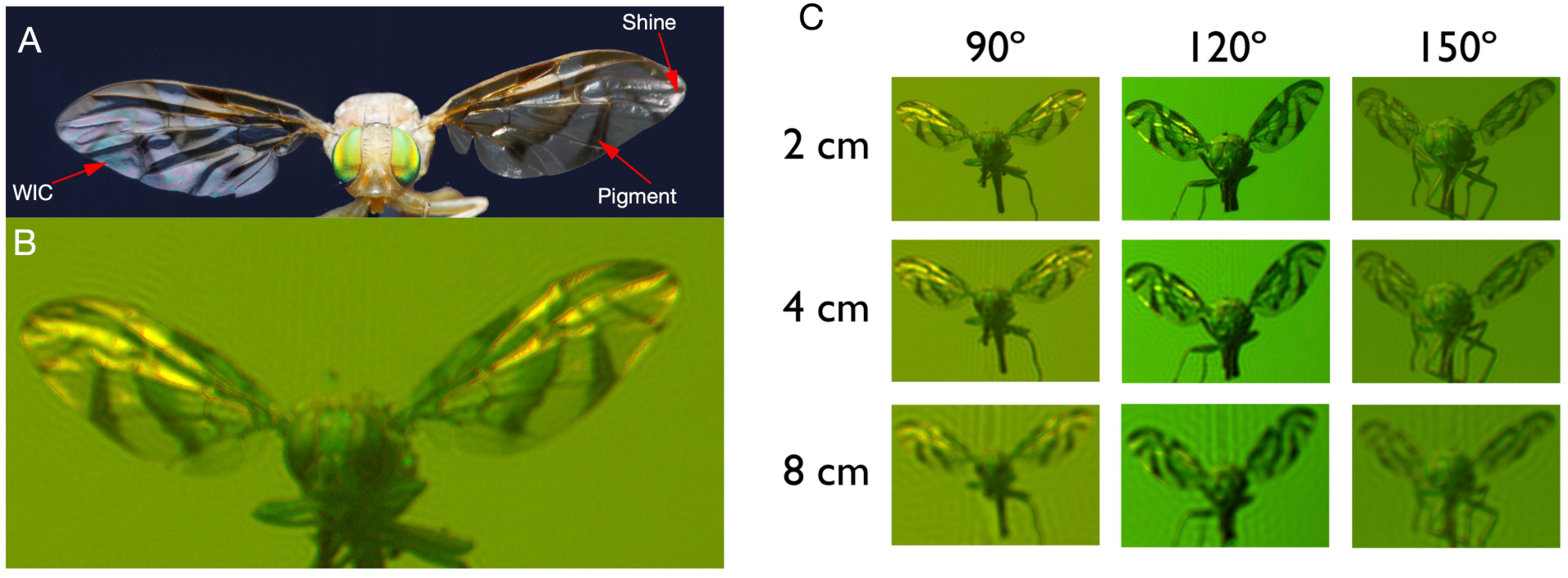
Simulation of the appearance of a Mexican fruit fly (*Anastrepha ludens*) as seen from the perspective of a jumping spider (A) Composite view of visual features showing wing specularity, wing interference colours and pigmentation. (B) Pseudo-colour image of the fly seen from the perspective of a *Phidippus audax* jumping spider’s visual system at a distance of 2 cm; (C) Pseudo-colour images of different wing angles and distances.

**Figure 6.**
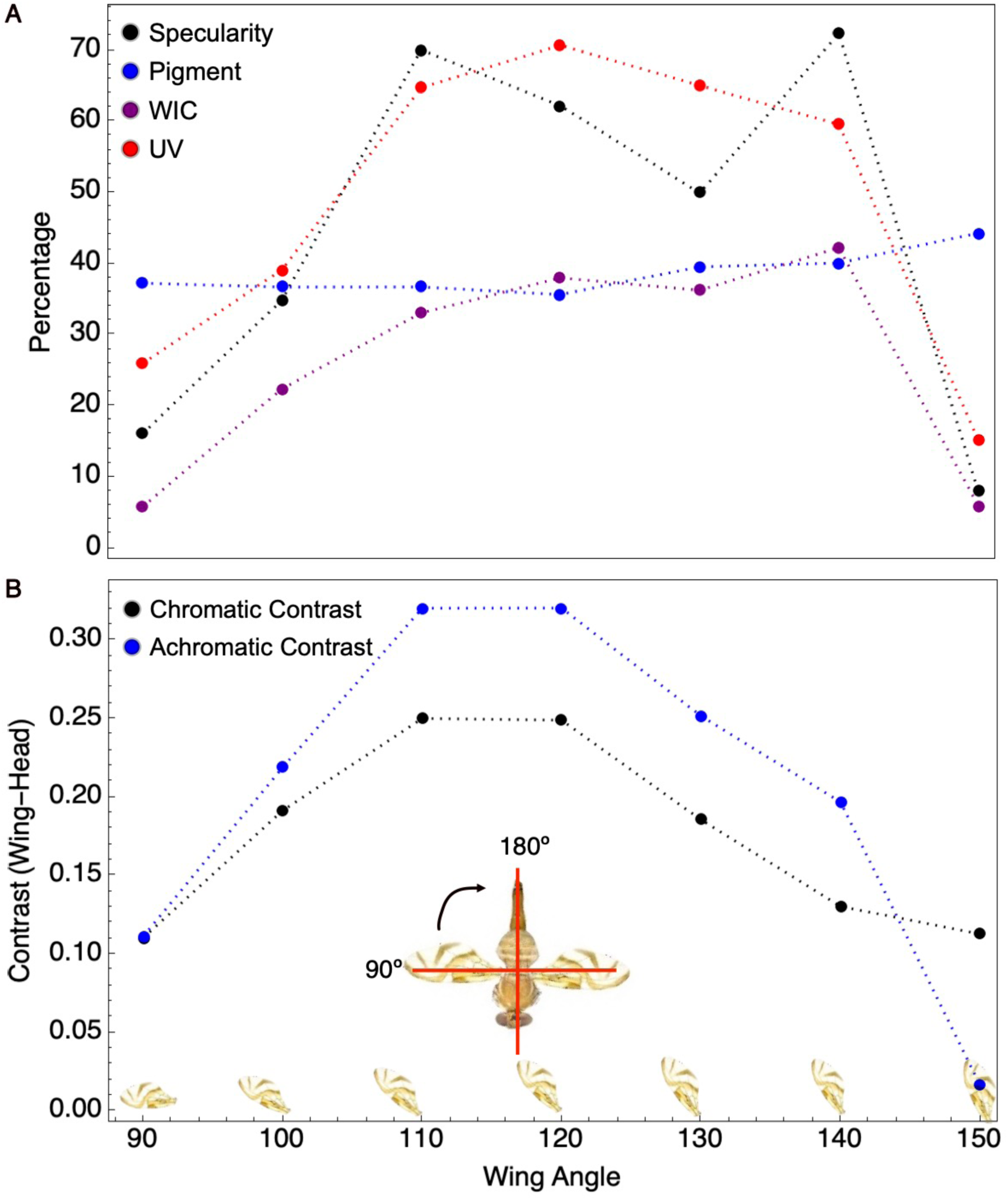
Appearance of fly’s wing feature from a spider’s perspective. (A) Apparent change in relative area of Pigment, Specularity, UV and WIC with opening angle of one wing; (B) Difference in achromatic and chromatic contrasts between the wing and the head with respect to opening angle of one wing. The contrast is highest when the wings are at the midpoint of display (110 -120°). Note that the dotted lines between the points are for visual clarity.

##### Achromatic and chromatic contrasts

The achromatic contrast (difference in luminance between wing and head) varied significantly according to the wing angle (Polynomial fit: R^2^ = 0.97, F = 95.54, p < 0.0005). The contrast followed a hump-shaped pattern, with a peak at 110-120° (Fig. 6B). A similar pattern was seen with the chromatic contrasts (Polynomial fit: R^2^ = 0.95, F = 23.69, p < 0.05; Fig. 6B). These results suggest that visual information as perceived by the spider fluctuates nonlinearly during the wing display.

##### Behavioural Assays

There was a significant difference in the percentage of flies that were attacked in different light conditions (Generalized Linear Regression; Logit Link function, Binomial distribution, χ^2^ = 8.09, df = 3, p < 0.044). With respect to the control, flies in the WIC with UV and WIC without UV treatments were attacked significantly less (Post Hoc contrasts; z = 2.11, p < 0.05 and z = 2.33, p < 0.05 respectively), while there was no significant difference between the Specularity treatment and the control (z = -1.2, p = 0.23; see S5 for the complete model).

## Discussion

In this paper, we bring together three different lines of evidence that suggests that the tephritid fly’s wing display generated a multitude of motion and colour cues which contributes to the hesitation by jumping spiders to attack. Using data from orientation in untethered spiders, retinal tracking in tethered spiders and visual modelling of wing appearance, we have presented a comprehensive picture of a deterrent visual display. The deterrent effect of a tephritid fly’s display against a jumping spider predator has been previously attributed to the mimetic markings on the fly’s wings (Greene et al., 1987). According to this hypothesis, spiders misidentify flies as other salticid spiders. However, salticids can be deterred even when the wing markings are artificially blackened out, suggesting that the wing motion has an important deterrent effect (Rao and Díaz-Fleischer, 2012). Our results provide a mechanism that can explain this deterrence. The hesitation to attack may be attributed to the failure by the spider to accurately track the movements of the fly. In the arena experiments, there is a fluctuation in the spider’s orientation when the fly is displaying, while the retina analysis showed that there was a significant lack of precision in tracking the motion of the fly. These subtle shifts in attention could cause the spider to slow or break off its attack, thereby allowing a displaying fly to survive an encounter with a predator (Rao and Díaz-Fleischer, 2012). In addition, the fly’s head does not move as much as the ovipositor and wings when the fly is displaying, perhaps to reduce the chances of the head to be detected and to attract attention to other, less vital, components. This behaviour is consistent with studies on other jumping spider species that use the head component of potential prey to target attacks (Bartos and Minias, 2016). From the fly’s point of view, reducing the motion of its head could allow the fly to gather more information about the nature of the threat.

To quantify the potential motion cues available from the fly movements, we calculated the optical flow of the movement from representative videos (Supplementary material S3, S4). The resulting vector field from the optical flow shows stark differences in magnitude and direction between a displaying fly and a moving fly. A displaying fly generates motion components in various directions and magnitudes which may make it more difficult for the spider to quickly identify the fly as prey. A moving fly, on the other hand, generates a uniform and single direction flow which potentially makes it easier for the spider to assess the best attack angle.

The hyaline parts of wings have been recently shown to produce bright structural colouration, and it has been suggested that predators may be dissuaded from attacking insects sporting such colours (Pike, 2015). In the jumping spider-tephritid system, there are special features that we need to consider. Firstly, jumping spiders are very sensitive to movement (Drees, 1952); their principal eyes have colour vision, but their anterior lateral eyes are ‘motion detection’ eyes (Jakob et al., 2018). Furthermore, orientation behaviour in salticids has been shown to be mediated by the anterior lateral eyes and is influenced by the speed and size of the stimuli (Zurek et al., 2010). The supination display of the fly produces a series of motion information: the wings themselves, the changing contrasts in multiple chromatic and achromatic perspectives as the wings are extended, and the movement of the head and legs as the display is performed. Note that since we only used one wing for the visual modelling, the number of colour and movement cues produced by the fly display is likely to be doubled under natural conditions.

The behavioural assays showed that spiders are less likely to attack flies in the wing interference colour treatment though they are likely to use only achromatic information in targeting prey. We detected no effect of UV on the likelihood of attacks by spiders. Since these spiders are dichromatic, with spectral sensitivity in the UV and green wavelengths, we suggest that only achromatic contrast is being used to target prey, i.e., the spiders possibly see the colour patterns as a ‘grayscale’ gradient (Voe, 1975). A similar effect may be operating in the red jumping spider *Saitis barbipes* that does not possess a red receptor (Glenszczyk et al., 2022).

Our results may be extended to other forms of displays. Male jumping spiders use elaborate courtship displays with an abundance of motion components, and mating success is correlated with the complexity of courtship display in at least some groups (Girard et al., 2015). Given that courtship can be dangerous for males, it may be that some complex visual displays are difficult for females to quickly assess, and thus buy time for a courting male to approach. These ideas need to be tested further.

## Acknowledgements

We thank Pablo Núñez Berea for help in collection of spiders in Mexico. Mary Emma Searles and Ashley Carey helped in the collection of retina data at UMass. We thank Nathan Morehouse and David Outomuro for helpful discussions. The eyetracker was designed by Cristina Canavesi and Jannick Rolland (University of Rochester) and built by Stingray Optics (Keene, North Hampshire). This project was financed by a CONACyT Mexico Ciencia Básica grant (No 168746) to DR, an NSF grant (IOS 0952822 and 1656714) to EJ and an Australian Research Council grant to AN (FT140100221, DP150101172)

## Author Contributions

Conceptualisation DR, EJ, SL

Formal Analysis DR, AN, HT-M, SL

Funding Acquisition DR, EJ

Investigation DR, KS-E, SA-A, LR-O, SL

Methodology DR, EJ, SL, LR-O

Project Administration DR, DR-M

Supervision DR, DR-M

Visualisation DR, HT-M, AN

Writing -original draft DR

Writing -review and editing DR, SL, EJ, AN

## Supplementary files

S1. Video showing supination display of the Mexican fruit fly (*Anastrepha ludens*) against a jumping spider predator (*Phidippus audax*).

S2. Sample video showing the response of the spider retinae overlaid with that of a displaying fly

S3. Results of a Kolomogorov Smirnov test looking at the distribution of the distance between the spider retinal midpoint and the stimulus (head of the fly) in the y axis (vertical displacement).

S4. Video showing pixel displacement of a displaying fly. The videos are colour coded according to magnitude of displacement, with redder pixels showing higher displacement and purple pixels showing lower displacement. See Fig. 4 for a vector field representation of these videos.

S5. Video showing pixel displacement of a moving fly. The videos are colour coded according to magnitude of displacement, with redder pixels showing higher displacement and purple pixels showing lower displacement. See Fig. 4 for a vector field representation of these videos.

S6. Generalized linear model looking at the effect of different light environments and wing appearance on predator attack

S**7**. Calculation of the minimum resolvable angle

## References

Aguilar-Argüello, S., Díaz-Fleischer, F. and Rao, D. (2015). Target-invariant aggressive display in a tephritid fly. Behavioural Processes 121, 33–36.

Aluja, M., Piñero, J., Jácome, I., Diaz-Fleischer, F. and Sivinski, J. (2006). Behavior of flies in the genus Anastrepha. In Fruit Flies (Tephritidae): Phylogeny and Evolution of Behavior (ed. Aluja, M.) and Norrbom, A.L.), pp. 376–401.

Bartos, M. and Minias, P. (2016). Visual cues used in directing predatory strikes by the jumping spider Yllenus arenarius (Araneae, Salticidae). Animal Behaviour 120, 51–59.

Brodie, E. D. (1977). Salamander Antipredator Postures. Copeia 1977, 523–535.

Canavesi, C., Long, S., Fantone, D., Jakob, E., Jackson, R. R., Harland, D. and Rolland, J. P. (2011). Design of a retinal tracking system for jumping spiders. In SPIE OPTICAL ENGINEERING + APPLICATIONS 2011 (ed. Koshel, R.J.) and Gregory, G. G.), pp. 81290A–8.

Carducci, J. and Jakob, E. (2000). Rearing environment affects behaviour of jumping spiders. Animal Behaviour (United Kingdom) 59, 39–46.

Caro, T. (1995). Pursuit-deterrence revisited. Trends in Ecology and Evolution 10, 500–503.

Caves, E. M. and Johnsen, S. (2017). AcuityView: An r package for portraying the effects of visual acuity on scenes observed by an animal. Methods Ecol Evol 9, 793–797.

Caves, E. M., Brandley, N. C. and Johnsen, S. (2018). Visual Acuity and the Evolution of Signals. Trends in Ecology and Evolution 33, 1–15.

Cronin, T. W., Johnsen, S., Marshall, N. J. and Warrant, E. J. (2014). Visual Ecology. Princeton University Press.

Drees, O. (1952). Untersuchungen über die angeborenen Verhaltensweisen bei Springspinnen (Salticidae). Zeitschrift für Tierpsychologie 9, 169–207.

Eichorn, C., Hrabar, M., Ryn, E. C. V., Brodie, B. S., Blake, A. J. and Gries, G. (2017). How flies are flirting on the fly. BMC Biology 15, 1–10.

Endler, J. A. and Mielke, P. W. (2005). Comparing entire colour patterns as birds see them. Biological Journal of the Linnean Society 86, 405–431.

Girard, M. B., Elias, D. O. and Kasumovic, M. M. (2015). Female preference for multi-modal courtship: multiple signals are important for male mating success in peacock spiders. Proceedings of the Royal Society of London. Series B: Biological Sciences 282, 20152222.

Glenszczyk, M., Outomuro, D., Gregorič, M., Kralj-Fišer, S., Schneider, J. M., Nilsson, D.-E., Morehouse, N. I. and Tedore, C. (2022). The jumping spider Saitis barbipes lacks a red photoreceptor to see its own sexually dimorphic red coloration. Sci Nat 109, 6.

Gonzalez-Lopez, G. I., Rao, D., Diaz-Fleischer, F., Orozco-Dávila, D. and Perez-Staples, D. (2015). Antipredator behavior of the new mass-reared unisexual strain of the Mexican Fruit Fly. Bulletin of Entomological Research 106, 314–321.

Greene, E., Orsak, L. J. and Whitman, D. W. (1987). A Tephritid Fly Mimics the Territorial Displays of Its Jumping Spider Predators. Science 236, 310–312.

Harland, D. P., Li, D. and Jackson, R. R. (2012). How Jumping Spiders See the World. In How animals see the world: Comparative behavior, biology, and evolution of vision (ed. Lazareva, O.F.), Shimizu, T.), and Wasserman, & E.A.), pp. 133–163.

Hasson, O. (1991). Pursuit-deterrent signals: Communication between prey and predator. Trends in Ecology and Evolution 6, 325–329.

Hasson, O. (1995). A Fly in Spiders Clothing - What Size the Spider. Proceedings of the Royal Society of London. Series B: Biological Sciences 261, 223–226.

Hawkes, M. F., Duffy, E., Joag, R., Skeats, A., Radwan, J., Wedell, N., Sharma, M. D., Hosken, D. J. and Troscianko, J. (2019). Sexual selection drives the evolution of male wing interference patterns. Proceedings of the Royal Society of London. Series B: Biological Sciences 286, 20182850–8.

Hill, D. E. (2022). Neurobiology and vision of jumping spiders (Araneae: Salticidae). Invertebr Syst 255.1, 1–81.

Jakob, E. M., Long, S. M., Harland, D. P., Jackson, R. R., Carey, A., Searles, M. E., Porter, A. H., Canavesi, C. and Rolland, J. P. (2018). Lateral eyes direct principal eyes as jumping spiders track objects. Current Biology 28, R1092–R1093.

Kelley, J. L., Tatarnic, N. J., Schröder-Turk, G. E., Endler, J. A. and Wilts, B. D. (2019). A Dynamic Optical Signal in a Nocturnal Moth. Current Biology 29, 2919-2925.e2.

Kjernsmo, K., Whitney, H. M., Scott-Samuel, N. E., Hall, J. R., Knowles, H., Talas, L. and Cuthill, I. C. (2020). Iridescence as Camouflage. Curr Biol 30, 551–555.

Land, M. (1969). Movements of the retinae of jumping spiders (Salticidae: Dendryphantinae) in response to visual stimuli. Journal of experimental biology 51, 471–493.

Ligon, R. A., Diaz, C. D., Morano, J. L., Troscianko, J., Stevens, M., Moskeland, A., Laman, T. G. and Scholes, E. (2018). Evolution of correlated complexity in the radically different courtship signals of birds-of-paradise. PLoS Biology 16, e2006962–24.

Mather, M. H. and Roitberg, B. D. (1987). A Sheep in Wolfs Clothing - Tephritid Flies Mimic Spider Predators. Science 236, 308–310.

Meijering, E., Dzyubachyk, O. and Smal, I. (2012). Methods for Cell and Particle Tracking. Methods in Enzymology 504, 183–200.

Morehouse, N. I. (2020). Spider vision. Current Biology 30, R975–R980.

Olsson, P., Lind, O., Kelber, A. and Simmons, L. (2018). Chromatic and achromatic vision: parameter choice and limitations for reliable model predictions. Behavioral Ecology 29, 273–282.

Otto, J. C. and Hill, D. E. (2017). Catalogue of the Australian peacock spiders (Araneae: Salticidae: Euophryini: Maratus, Saratus). Peckhamia 1–21.

Parker, A. (1998). The diversity and implications of animal structural colours. Journal of experimental biology 201, 2343–2347.

Peaslee, A. G. and Wilson, G. (1989). Spectral sensitivity in jumping spiders (Araneae, Salticidae). Journal of Comparative Physiology, A 164, 359–363.

Pike, T. W. (2015). Interference coloration as an anti-predator defence. Biology Letters 11, 20150159–20150159.

Rao, D. and Díaz-Fleischer, F. (2012). Characterisation of Predator-Directed Displays in Tephritid Flies. Ethology 118, 1165–1172.

Raudies, F. (2013). Optic flow. Scholarpedia 8, 30724.

Robledo-Ospina, L. E., Escobar-Sarria, F., Troscianko, J. and Rao, D. (2017). Two ways to hide: predator and prey perspectives of disruptive coloration and background matching in jumping spiders. Biol J Linn Soc 122, 752–764.

Rosenthal, G. G. (2007). Spatiotemporal Dimensions of Visual Signals in Animal Communication. Annual Review of Ecology, Evolution, and Systematics 38, 155–178.

Rößler, D. C., Agrò, M. D., Kim, K. and Shamble, P. S. (2021). Static visual predator recognition in jumping spiders. Funct Ecol.

Ruxton, G., Sherratt, T. N. and Speed, M. (2018). Avoiding Attack: the evolutionary ecology of crypsis, warning signals and mimicry. 2nd ed. Oxford Univerity Press.

Schultz, T. D. and Fincke, O. M. (2009). Structural colours create a flashing cue for sexual recognition and male quality in a Neotropical giant damselfly. Functional ecology(Print) 23, 724–732.

Shevtsova, E., Hansson, C., Janzen, D. H. and Kjaerandsen, J. (2011). Stable structural color patterns displayed on transparent insect wings. Proceedings of the National Academy of Sciences 108, 668–673.

Stuart-Fox, D., Ospina-Rozo, L., Ng, L. and Franklin, A. M. (2020). The Paradox of Iridescent Signals. Trends in Ecology and Evolution 6, 1–9.

Troscianko, J. and Stevens, M. (2015). Image calibration and analysis toolbox – a free software suite for objectively measuring reflectance, colour and pattern. Methods Ecol Evol 6, 1320–1331.

Voe, R. D. D. (1975). Ultraviolet and green receptors in principal eyes of jumping spiders. The Journal of General Physiology 66, 193–207.

Vorobyev, M. and Osorio, D. (1998). Receptor noise as a determinant of colour thresholds. Proc Royal Soc Lond Ser B Biological Sci 265, 351–358.

Winsor, A. M., Pagoti, G. F., Daye, D. J., Cheries, E. W., Cave, K. R. and Jakob, E. M. (2021). What gaze direction can tell us about cognitive processes in invertebrates. Biochemical and Biophysical Research Communications 87, 39–12.

Zurek, D. B., Taylor, A. J., Evans, C. S. and Nelson, X. J. (2010). The role of the anterior lateral eyes in the vision-based behaviour of jumping spiders. Journal of Experimental Zoology Part A: Ecological Genetics and Physiology 213, 2372–2378.

